# Rapid phenotypic and metabolomic domestication of wild *Penicillium* molds on cheese

**DOI:** 10.1101/647172

**Authors:** Ina Bodinaku, Jason Shaffer, Allison B. Connors, Jacob L. Steenwyk, Erik Kastman, Antonis Rokas, Albert Robbat, Benjamin Wolfe

**Affiliations:** Tufts University, Department of Biology, Medford, MA, USA; Tufts University, Department of Chemistry, Medford, MA, USA; Vanderbilt University, Department of Biological Sciences, Nashville, TN, USA; Tufts University Sensory and Science Center, Medford, MA, USA

## Abstract

Fermented foods provide novel ecological opportunities for natural populations of microbes to evolve through successive recolonization of resource-rich substrates. Comparative genomic data have reconstructed the evolutionary histories of microbes adapted to food environments, but experimental studies directly demonstrating the process of domestication are lacking for most fermented food microbes. Here we show that during the repeated colonization of cheese, phenotypic and metabolomic traits of wild *Penicillium* molds rapidly change to produce mutants with properties similar to industrial cultures used to make Camembert and other bloomy rind cheeses. Over a period of just a few weeks, populations of wild *Penicillium* strains serially passaged on cheese resulted in the reduction or complete loss of pigment, spore, and mycotoxin production. Mutants also had a striking change in volatile metabolite production, shifting from production of earthy or musty volatile compounds (e.g. geosmin) to fatty and cheesy volatiles (e.g. 2-nonanone, 2-undecanone). RNA-sequencing demonstrated a significant decrease in expression of 356 genes in domesticated mutants, with an enrichment of many secondary metabolite production pathways in these downregulated genes. By manipulating the presence of neighboring microbial species and overall resource availability, we demonstrate that the limited competition and high nutrient availability of the cheese environment promote rapid trait evolution of *Penicillium* molds.

**IMPORTANCE:** Industrial cultures of filamentous fungi are used to add unique aesthetics and flavors to cheeses and other microbial foods. How these microbes adapted to live in food environments is generally unknown as most microbial domestication is unintentional. Our work demonstrates that wild molds closely related to the starter culture *Penicillium camemberti* can readily lose undesirable traits and quickly shift toward producing desirable aroma compounds. In addition to experimentally demonstrating a putative domestication pathway for *P. camemberti*, our work suggests that wild *Penicillium* isolates could be rapidly domesticated to produce new flavors and aesthetics in fermented foods.

## INTRODUCTION

Fermented foods such as cheese, miso, sourdough, and sauerkraut are hybrid microbiomes where wild microbial species from the environment mix with domesticated microbes that are added as starter cultures. Microbes from natural ecosystems have the potential adapt to these resource-rich environments where they may rapidly evolve new traits and/or lose traits that are not maintained by selection in these novel environments (1). Previous evidence for microbial domestication in fermented foods comes from comparative genomic studies of food isolates and closely-related wild strains (1–4). For example, in the fungus *Aspergillus oryzae*, which is used in the production of soy sauce, miso, and sake, both structural and regulatory genomic changes are correlated with the evolution of non-toxic and flavorful *Aspergillus oryzae* strains from a highly toxic ancestor (*Aspergillus flavus*) (2). The evolutionary origins of most domesticated microbes remain enigmatic in large part because domestication of microbes is usually unintentional and the processes driving microbial domestication have not been experimentally recreated (1).

*Penicillium* species colonize the surfaces of aged cheeses around the world either as starter cultures that are intentionally added during the cheese making process (5, 6) or as non-starter *Penicillium* species that enter cheese production facilities from natural fungal populations (7–9). The white surface of Camembert and Brie cheeses is created by a variety of strains of the starter culture *P. camemberti.* This domesticated fungus is white, makes fewer conidia (asexual spores) than most wild *Penicillium* species, does not make detectable levels of mycotoxins, and produces desirable mushroomy and fatty volatiles during cheese ripening (10–12) (**Fig. 1A**). In contrast, the putative ancestor *P. commune* and other closely related *Penicillium* species are generally greenish-blue, make large numbers of conidia, and produce mycotoxins and other undesirable volatiles that negatively impact cheese quality (**Fig. 1B-C**). Historical accounts suggest that white mutant strains of *Penicillium* species were either directly isolated from French cheeses or produced in laboratories in France (13), but the potential domestication processes that generated these iconic cheese mold species have not been identified.

**Figure 1:**
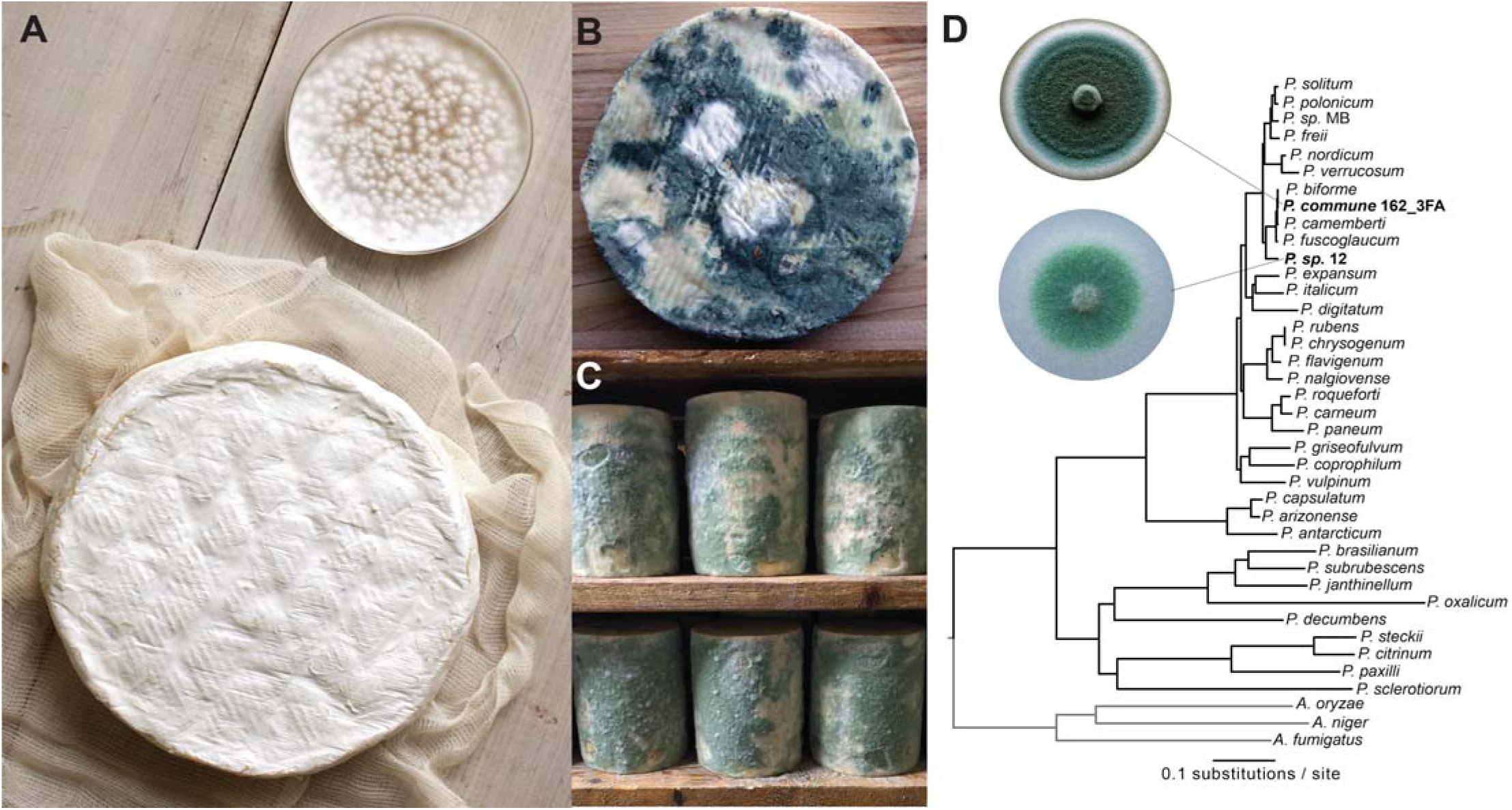
*Penicillium* molds in the cheese environment. **(A)** The white mold known as *Penicillium camemberti* (show in pure culture in the Petri dish) is used to make Camembert (shown), Brie, and other bloomy rind cheeses. **(B)** Wild *Penicillium* molds (also known as non-starter molds) can contaminate cheeses during production. **(C)** Some natural rind cheeses are intentionally colonized by wild *Penicillium* molds. Shown here is *Penicillium* sp. strain 12, a strain used in the experiments in this paper, colonizing wheels of a blue cheese in a cave in the United States. **(D)** A phylogenomic tree of *Penicillium*. Strains used in this work are highlighted in bold. *Penicillium* sp. MB was also isolated from a natural rind cheese and sequenced as part of this work, but was not used in the experiments described.

Here we used experimental evolution to determine how wild *Penicillium* molds may be unintentionally domesticated in the cheese aging environment. We specifically determined how quickly *Penicillium* could evolve new phenotypes on cheese, how *Penicillium* traits change during domestication on cheese, and what properties of the cheese environment promote domestication of *Penicillium*. Using our cheese rind model (14, 15), we serially passaged populations of wild *Penicillium* and tracked the emergence of phenotypic mutants. We found that mutants with substantially reduced mycotoxin levels, reproductive output, and pigment production rapidly emerge in these experimental populations. Volatile profiling and RNA-sequencing also demonstrate a substantial remodelling of metabolism in domesticated mutants. These findings illustrate the potential for rapid domestication of filamentous fungi in cheese caves around the world.

## RESULTS and DISCUSSION

### Non-starter *Penicillium* species rapidly evolve novel phenotypes on cheese

To experimentally evolve *Penicillium* on cheese, two non-starter *Penicillium* strains (*Penicillium commune* strain 162_3FA and *Penicillium* sp. strain 12) isolated from a cheese cave in Vermont in the United States were serially passaged on cheese curd medium. These molds were isolated from a cheese aging facility that was colonized by the molds within the past five years. These cheese cave isolates have wild-type phenotypes (pigmented, high spore production, musty odors, and mycotoxin production), and are closely related to *P. camemberti* strains used in cheese production (**Fig. 1D**, **Fig. S1**). At each passage, replicate populations were sampled to determine population size and mutant frequency. Colonies were considered phenotypic mutants if they had altered surface color or texture indicating changes in pigment or spore production. To determine how competition from neighboring cheese microbes impacts the rate of phenotypic diversification, we serially passaged replicate populations alone (“*Penicillium* alone”) or in the presence of a mix of three competitors (“*Penicillium* + community”, including the yeast *Debaryomyces hansenii*, and the bacteria *Brachybacterium alimentarium* and *Staphylococcus xylosus*) that commonly co-occur with *Penicillium* species in cheese rinds (14, 16, 17).

Within four weeks of serial passage on cheese, mutant phenotypes began to emerge in our experimental *Penicillium* populations, reaching 71.5% of the population in the *Penicillium* alone treatments by the end of the experiment (**Fig. 2A**). The presence of neighbors strongly inhibited mutant phenotype frequency in the *Penicillium* + community treatments (mean of 26.2%, repeated-measures ANOVA *F*_1,6_= 86.5, *p* < 0.001, Fig. 2A). Neighbors also decreased total population size with an average of 42% decrease in total CFUs across the experiment (repeated-measures ANOVA *F*_1,6_= 10.3, *p* = 0.02, **Fig. S1**). Similar patterns of phenotypic diversification alone and inhibition with neighbors were observed with *Penicillium* sp. strain 12 (**Fig. S2A-B**, **Table S2A-S2B**). These results suggest that cheese can promote the rapid phenotypic diversification of *Penicillium* molds and that biotic interactions in cheese rinds can inhibit this diversification.

**Figure 2:**
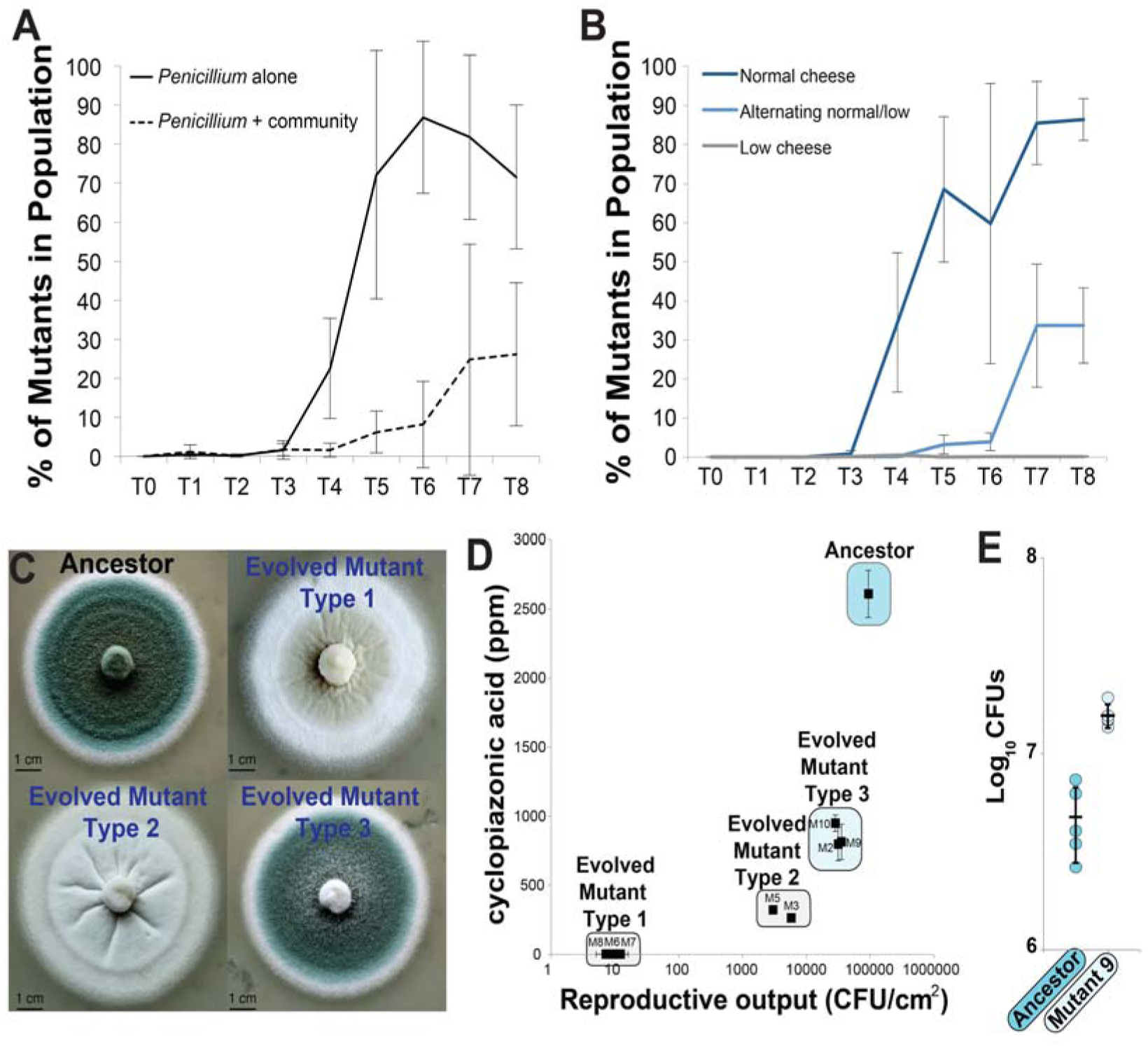
Experimental evolution of *Penicillium* on cheese. **(A)** Evolution of *Penicillium commune* strain 162_3FA on cheese curd alone (“*Penicillium* alone”) and in the presence of three competing cheese rind microbes (“*Penicillium* + community”; *Staphylococcus xylosus, Brachybacterium alimentarium,* and *Debaryomyces hansenii*). Lines connect points representing mean mutant phenotype frequencies of four replicate populations and error bars represent one standard deviation of the mean. “*Penicillium* + community” had a significantly lower mutant frequency (repeated-measures ANOVA, see text for stats). **(B)** Experimental evolution of *P. commune* strain 162_3FA in different cheese nutrient environments. “Normal cheese” = 10% cheese curd in agar medium. “Low cheese” = 1% cheese curd in agar medium. “Alternating normal/low” = alternating 10% and 1% cheese curd at each transfer. Both “Low cheese” and “Alternating normal/low” had significantly lower mutant frequencies (repeated-measures ANOVA with Tukey’s HSD post-hoc tests, see text for stats). Lines connect points representing mean mutant phenotype frequencies of four replicate populations and error bars represent one standard deviation of the mean. The “low cheese” line is difficult to see because it is at 0%. **(C)** Morphology of four representative colony types. The Ancestor wild-type phenotype which is dark blue-green and dusty, Evolved Mutant Type 1 which was white and flat, Evolved Mutant Type 2 which was white and fuzzy/dusty, and Evolved Mutant Type 3 which was blue-green, but had less intense coloration than wild-type and less fuzzy appearance. **(D)** Reproductive output and cyclopiazonic acid production of a range of mutants isolated across the experimental evolution populations. Points are mean values and error bars are one standard deviation of the mean. M5, M6, M7, and M8 had reduced reproductive compared to the Ancestor (Dunnett’s test, *p* < 0.05). All mutants had significantly reduced CPA production compared to the Ancestor (Dunnett’s test, *p* < 0.05). **(E)** Competition between the wild-type Ancestor *P. commune* strain 162_3FA and M9. M9 outcompeted the Ancestor after 10 days of growth on cheese curd agar. Points represent individual replicate competition communities and horizontal line indicates mean values. Error bars are one standard deviation.

Cheese is a high-resource environment compared to environments where *Penicillium* molds naturally occur (soils, leaves, etc.), with an abundance of carbon, nitrogen, and other resources stored in the protein casein (18). To determine if the high resource availability of cheese promotes the rapid domestication of *Penicillium* molds, we repeated the short-term experimental evolution above with the standard cheese curd (“normal cheese”), with a “low cheese” treatment with 1/10th the normal amount of cheese curd, and with an “alternating cheese” treatment with alternating normal and 1/10th cheese curd agar at every other passage. The low cheese treatment was designed to reduce total nutrient availability while keeping pH and other environmental variables similar. The alternating treatment was designed to simulate alternating colonization of a high resource environment (cheese) and a low resource environment (soil, wood) which could occur in a cheese aging facility.

As with our first set of experiments, the normal cheese treatment resulted in the rapid evolution of phenotypic mutants by the fourth week of the experiment with a mean mutant frequency of 89.0% (+/- 12.2) across four replication populations at the end of the experiment (**Fig. 2B**). Both the alternating cheese and low cheese treatment had significantly lower mutant frequencies across the duration of the experiment (mean mutant frequency at end of experiment: alternating cheese = 73.0 +/- 14.6%, low cheese= 0.16 +/- 0.19%; repeated-measures ANOVA *F*_2,9_= 149.6, *p* < 0.0001). As with the competition treatment above, population sizes were significantly lower in the low cheese treatment (repeated-measures ANOVA *F*_2,9_= 105.1, *p* <0.0001) and may explain the substantially suppressed rate of diversification (**Fig. S4**). These results suggest that the high resource environment of cheese promotes the rapid trait evolution of *Penicillium*.

### Domestication of *Penicillium* on cheese leads to stable reductions in reproductive output, pigmentation, and mycotoxin production

In our work above, all strains that emerged with altered colony morphotypes were grouped together to obtain overall rates of phenotypic evolution in different biotic and abiotic environments. To provide a finer-scale analysis of how reproductive and metabolic traits shifted during adaptation to cheese, we measured reproductive output and mycotoxin production of representative strains of *P. commune* 162_3FA that spanned the spectrum of mutant colony phenotypes (**Tables S1-S2**). Reproductive output was measured as the number of CFUs produced per unit area of a fungal colony and measured both spore and hyphal propagules. We also measured production of the mycotoxin cyclopiazonic acid (CPA) by ancestral and mutant strains when grown on cheese curd. Cyclopiazonic acid is commonly produced by *Penicillium commune* and other closely related *Penicillium* species that colonize cheese surfaces, but it is generally not produced or only produced in small quantities by *P. camemberti* strains used in cheese production (7, 12). Pigment production was also qualitatively described by photographing colonies of each strain grown on cheese curd agar. Spore, mycotoxin, and pigment production are traits that are often co-regulated by global regulators in *Aspergillus* and *Penicillium* species (19, 20). Mycotoxin and pigment production are thought to be important traits for filamentous fungi to compete with other microbes or tolerate oxidative stress (21, 22); however, these traits are frequently lost within fungal populations, suggesting that they are costly (23). We predicted that adaptation to cheese might lead to the loss of some of these costly traits in the high-resource and reduced competition environment of cheese rinds.

Trait analysis of the ancestor and seven mutant strains found a general pattern of reduced reproductive output, mycotoxin, and pigment production. Reproductive output was significantly lower in all mutant phenotypes compared to the ancestral strain (ANOVA with Dunnett’s test: *F*_8,18_=178.6, *p* < 0.001), with some strains having approximately 4-log reductions in spore production (**Fig. 2D**). We did not separate spores from hyphae in our analysis of reproductive output. However, we suspect the drop in reproductive output is largely due to loss of spore production because colonies of white mutants lost the characteristic dusty appearance created by spore-bearing conidiophores (**Fig. 2C**). We also detected a substantial loss of CPA production across all mutants of *P. commune* 162_3FA (ANOVA with Dunnett’s test: *F*_8,18_=38.2, *p* < 0.001), with three strains (M6, M7, and M8) having no detectable levels of CPA. Pigment production followed the pattern of loss of reproductive output and CPA production, with intermediate light blue mutants that had intermediate levels of reproductive output and CPA production (M2, M9, and M10) and completely white mutants having the lowest levels of reproductive output and CPA production (M3, M5, M6, M7, and M8). The observed phenotypic changes were not transient; repeated transfer of mutants on cheese curd agar did not lead to reversions to wild-type morphologies (**Fig. S5**). These trait analyses suggest co-regulated loss of reproductive output, mycotoxin production, and pigmentation during adaptation of *P. commune* to the cheese environment.

The fitness of mutants may differ from the ancestral strain when growing on the rich cheese medium because mutants may shift resource allocation from costly traits (e.g. secondary metabolite production) to growth. To test whether mutants had a higher fitness compared to the ancestor, we used competition experiments where equal amounts of a mutant strain (M9) were co-inoculated with the ancestor (WT). We were only able to conduct these competition experiments with mutants that still had some level of spore production because it is difficult to standardize inputs of wild-type spore producers and white mutant strains that had almost no spore production. As predicted, the mutant outcompeted the ancestor after ten days (**Fig. 2E**), suggesting that the coordinated loss of traits during domestication leads to a higher fitness in the cheese environment.

### Domestication shifts the *Penicillium* volatilome from musty to cheesy aromas

While working with the mutant strains, we noticed that they had strikingly different aromas compared to the wild-type ancestral strain. The ancestral strain smelled musty and earthy while the mutants smelled fatty, green, and surprisingly reminiscent of aged cheese. The aromas of cheese are volatile organic compounds (VOCs) that are produced by filamentous fungi and other microbes during proteolysis, lipolysis, and other processes that decompose the cheese substrate (11, 24, 25). These VOCs are important determinants of how consumers perceive the quality of a cheese and can create variation in aroma profiles across surface-ripened cheese varieties (26, 27). To quantitatively assess whether domestication of *Penicillium* on cheese alters volatile aroma production, we captured volatiles produced by wild-type and mutant *Penicillium* using headspace sorptive extraction followed by analysis with gas chromatography-mass spectrometry (GC-MS) (28, 29). We compared the ancestor *P. commune* 162_3FA with three mutants - M2, M5, and M6 - that spanned the continuum of reproductive output, mycotoxin, and pigment traits (**Fig. 2D**).

As suggested by our preliminary olfactory observations, the composition of volatiles produced by mutants shifted substantially from the ancestor (**Fig. 3**, ANOSIM *R* = 1.0, *p* < 0.001). Geosmin was the only VOC that was produced by the ancestor and was absent in all three of the mutant strains (6.722% contribution in SIMPER analysis of ancestor vs. all mutants pooled together; **Fig. 3**, **Table S3**). Geosmin is widely recognized for contributing a musty aroma to environmental and food samples and is produced by both bacteria and fungi including *Penicillium* (30–32). It has a very low odor threshold meaning that even small amounts of geosmin produced by *Penicillium* can be perceived as strong aromas (33). The loss of geosmin production could be the major driver of the perceived loss of musty aromas in domesticated *Penicillium* mutants.

**Figure 3:**
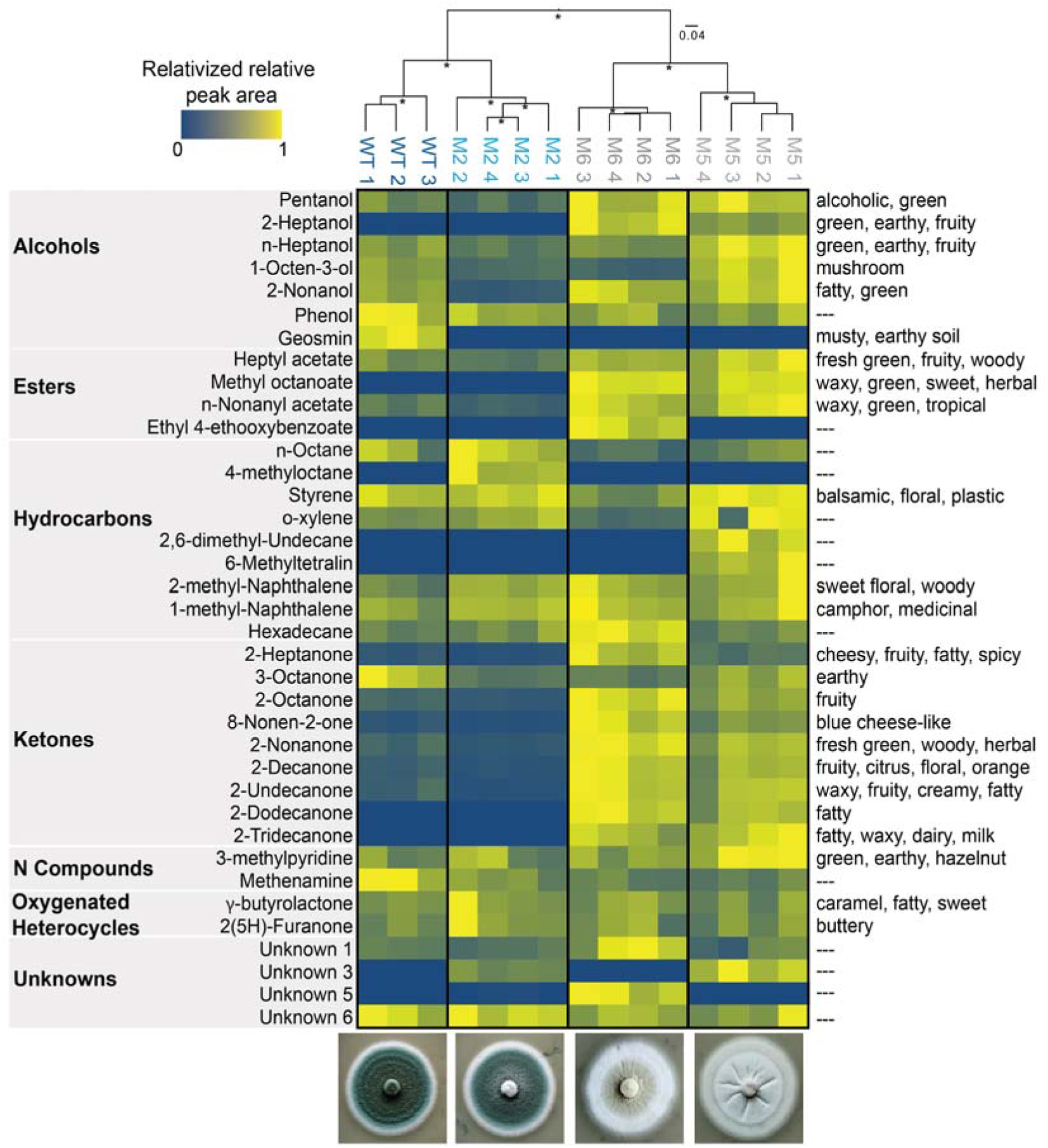
Volatile organic compound (VOC) production of wild-type and domesticated *Penicillium*. Because total concentrations of VOCs are highly variable across different compounds, visualization was simplified by relativizing the relative peak area from GC-MS chromatograms within each VOC to the highest concentration detected for that VOC. Only VOCs that were detected across all replicates are shown. See Table S3 for all VOCs and their relative peak area values. The UPGMA tree is clustering the VOC profiles for each replicate based on Bray-Curtis dissimilarity. Asterisks indicate clusters with > 70% bootstrap support. WT = ancestor wild-type. M2, M5, and M6 are all mutants. Numbers after strains (1,2,3,4) indicate biological replicates. Because of accidental sample loss during processing, only three biological replicates were collected from WT. Descriptors on the right are known aroma qualities of detected VOCs.

In addition to a loss of musty aromas, mutant strains produced higher amounts of methyl ketones and other VOCs associated with molds used in cheese production. Typical Camembert flavor has been defined in patents as containing 2-heptanone, 2-heptanol, 8-nonen-2-one, 1-octen-3-ol, 2-noanol, phenol, butanoic acid, and methyl cinnamate (11). All of these VOCs, except butanoic acid and methyl cinnamate, were detected in our GC-MS profiling and several were major drivers of differences in VOC profiles of mutants compared to the ancestor (**Fig. 3;** 2-heptanone = 12.1% SIMPER contribution, 8-nonen-2-one = 8%, 1-octen-3-ol = 7% contribution). Other methyl ketones that have been detected in *P. camemberti* (34), including 2-nonanone (13.8% contribution) and 2-undecanone (12.3%), were also detected in higher concentrations in mutants compared to the ancestor and contributed strongly to differences in VOC profiles (Fig. 3). These methyl ketones are perceived as cheesy, fatty, fruity, and green aromas that are typically associated with ripened cheeses (35, 36). Collectively, these VOC data demonstrate a dramatic remodelling of the volatilome of *P. commune* as a result of rapid domestication on cheese.

### Comparative transcriptomics demonstrates global down-regulation of secondary metabolite production in domesticated *Penicillium* mutants

To explore additional shifts in metabolic processes in cheese-adapted *Penicillium* not captured by our targeted metabolomics above, we used RNA-sequencing (RNA-seq) to compare global expression patterns of the wild-type *P. commune* to one mutant strain (M5). This mutant was selected because it had an intermediate reduction of reproductive output, CPA production, and pigment production. We predicted that in addition to shifts in gene expression related to spore and pigment production, genes associated with other secondary metabolites not measured would also be downregulated.

The transcriptome of mutant M5 had 356 genes with significantly lower expression compared to the ancestor, or about 3.2% of all protein-coding genes (**Fig. 4A**). Only 86 genes had higher expression levels in the mutant compared to the ancestor. An enrichment analysis of GO terms associated with these differentially expressed genes highlights the substantial downregulation of genes associated with secondary metabolite production (**Fig. 4B**). Many pathways that were significantly enriched in the list of downregulated genes were associated with pigment production (melanin biosynthesis) and production of a range of secondary metabolites including chanoclavine-I, austinol, and dehydroaustinol (**Fig. 4B**, **Table S4**). One striking example is the ergot alkaloid synthesis (*eas*) gene cluster. Ergot alkaloids can be toxic to mammals and recent work demonstrated that genomes of *P. camemberti* from cheese contain some genes in the ergot alkaloid biosynthesis pathway and can produce some early precursors of ergot alkaloids (37). The *eas* gene cluster is also present in *P. commune* 162_3FA and is strongly downregulated (a mean of −42 fold-change across *dmwA*, *easE*, *easF*, and *easC* genes) in the mutant compared to the ancestor (**Fig 4C**). In addition to the dramatic decrease in the expression of genes associated with secondary metabolite production, the observed reduction in conidia production by mutants is supported by strong downregulation of *abaA*, which regulates condia development in *Aspergillus* (38) (**Table S4**).

**Figure 4:**
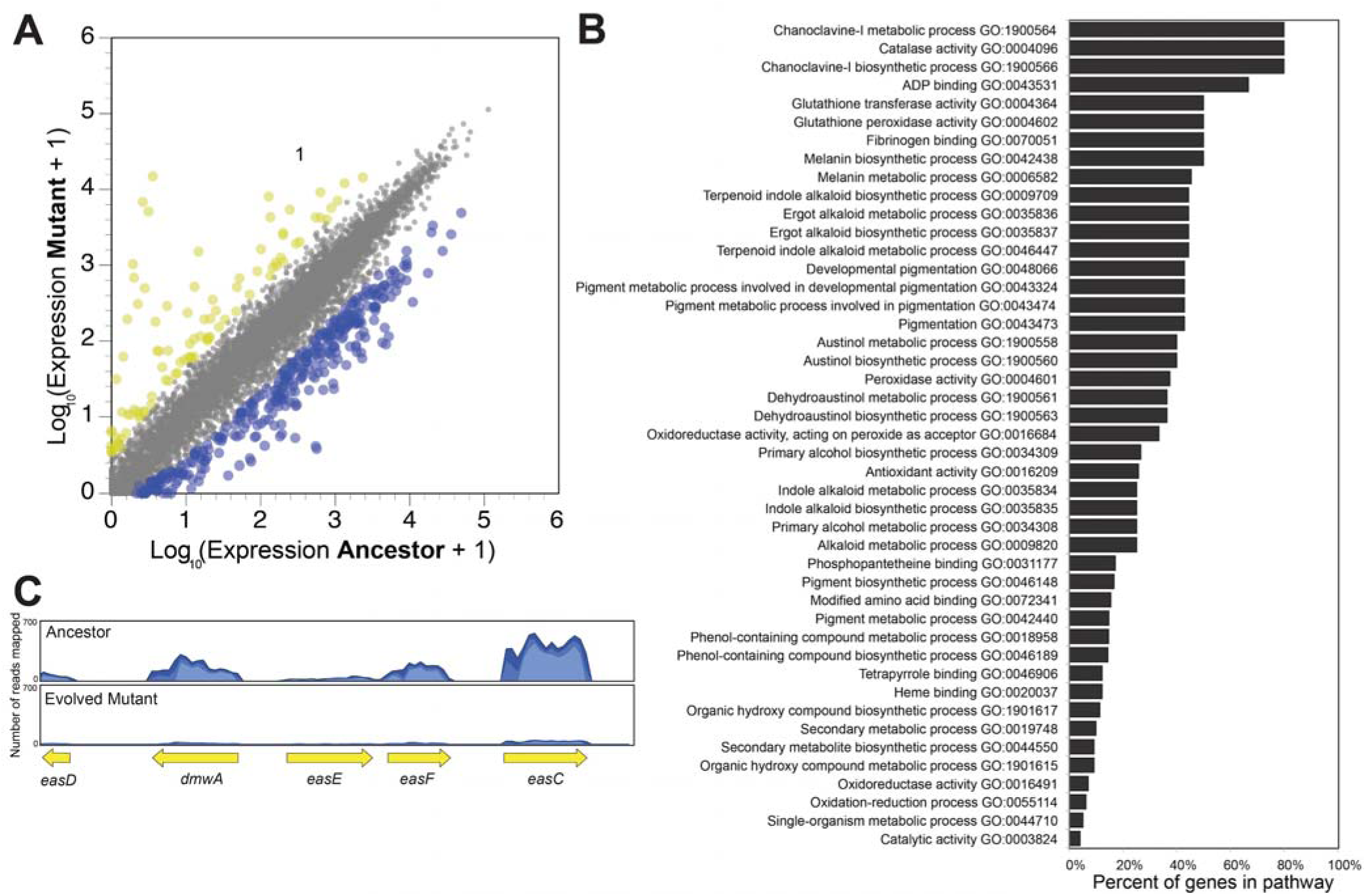
Experimental domestication shifts global gene expression of Penicillium on cheese. **(A)** Differences in gene expression between ancestor (wild-type) and mutant strain M5 *Penicillium commune* 162_3FA. Each dot represents a transcript from across the genome. Yellow dots represent those transcripts that had higher expression and blue dots represent those transcripts that had lower expression (5-fold change in expression, FDR corrected p-value < 0.05). **(B)** Pathway enrichment analysis showing distribution of GO terms that were significantly enriched in genes with decreased expression in Mutant (strain M5). **(C)** Representative mapping of reads to the ergot alkaloid synthesis (*eas*) gene cluster.

**Figure 5:**
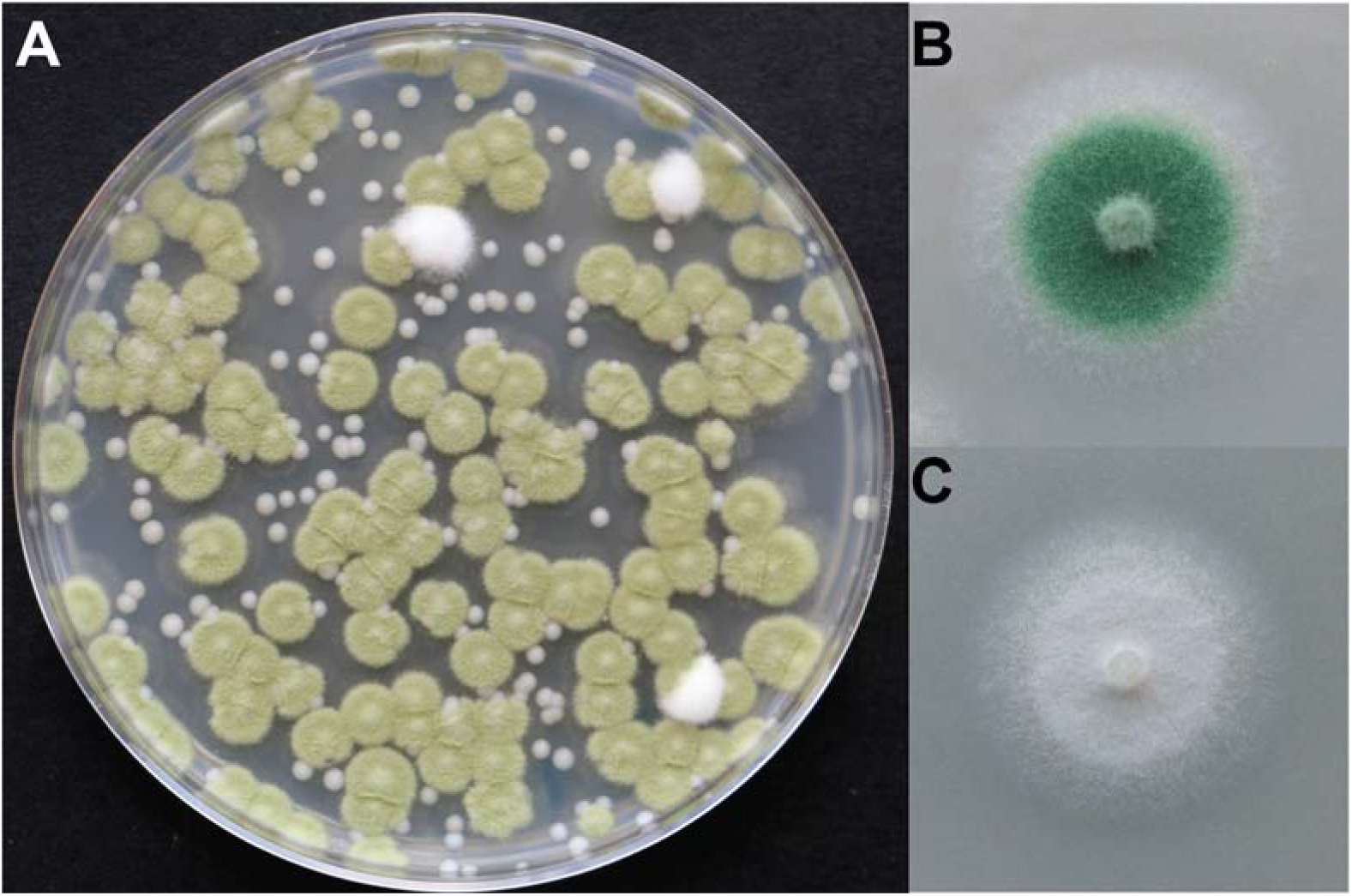
Phenotypic mutants of *Penicillium* are present in cheese caves. **(A)** A plate showing *Penicillium* sp. strain 12 isolated from from a cheese cave in Vermont, USA. The fuzzy white colonies are white mutants that exist at low frequencies in this fungal population. The small smooth beige colonies are the yeast *Debaryomyces hansenii*. **(B)** Wild-type phenotype of the mold isolated from the population and grown on cheese curd agar. Green pigmentation is different between A and B because A is grown on plate count agar and B is grown on cheese. **(C)** A white mutant isolated from the same population as the wild-type.

### Domesticated mutants of *Penicillium* are found at low frequencies in cheese caves

Our work above provides experimental evidence that *Penicillium* molds can rapidly domesticate in the cheese environment. But does this domestication produce phenotypic mutants in the more realistic conditions of a cheese cave? To answer this question, we deeply sampled a population of *Penicillium* sp. strain 12 from a cheese cave in Vermont, USA. Sampling of *Penicillium* sp. strain 12 occurred four years after the initial isolation of this strain. We removed patches of the fungus from the surface of 43 different wheels of cheese and plated out each of the patch samples to determine the frequency of wild-type vs. mutant colonies. Much of our experimental work focused on *P. commune* 162_3FA and ideally we would have sampled a population of this fungus. However, we were unable to find a large enough population of *P. commune* 162_3FA in the cave where it was originally isolated.

White mutant colonies were detected on 12 of the 43 wheels of cheese and were infrequent relative to wild-type (0.36% frequency). This low abundance of mutants in these multispecies rind communities aligns with the low mutant frequency observed when *Penicillium* sp. strain 12 evolved in the presence of competitors (2.23%, **Fig S3B**). This survey demonstrates that phenotypic mutants of wild molds can be detected in caves where cheeses are aged. These rare mutants were likely the source of the original white molds used in industrial Camembert production.

## Conclusions

Novel ecological opportunities are thought to promote the diversification of plants and animals during adaptive radiations (39–41) and similar processes may occur when wild microbial populations colonize the high-resource environments of fermented foods. Cheese is a resource-rich substrate that provides microbes from natural populations with novel ecological opportunities. Facilities where natural rind cheeses are aged are relatively stable environments where stressors of the natural world, including resource-limitation and UV stress, are relaxed. While these facilities are carefully managed to keep out pathogens, wild molds from natural populations commonly colonize the surfaces of certain cheeses where a natural rind is desired (6, 7, 14). Using experimental evolution, we demonstrate that cheese aging environments have the potential to promote rapid trait evolution of *Penicillium* species. Changes in the cheese environment that suppressed population size, including competition and decreased resource availability, inhibited trait evolution during domestication. Previous comparative genomic studies of *Penicillium* molds from cheese and fermented meat have identified genomic signatures of evolution over longer time-scales in fermented food environments (3, 42). Our experimental work demonstrates that in just a few weeks, *Penicillium* molds can adapt to the cheese environment through the loss of energetically costly traits, including spore production, pigment production, and mycotoxin production.

Previous studies in *Aspergillus* and *Penicillium* species have observed similar rapid trait change when fungi are subcultured in rich lab media over many generations (43–45). This phenomenon has been called degeneration because desired industrial traits or traits of interest for laboratory studies are lost. These cultures experience a similar transition from a high competition and low resource natural environment (plants, soil, etc.) to the low competition and high resource environment (rich lab media) as is the case when fungi colonize cheese. For example, serial transfer of *A. parasiticus* caused the rapid loss of secondary metabolite production, decreased sporulation, and changes in pigment production at a similar timescale (several weeks) to that observed in our work on cheese (44). A similar pattern of trait degeneration with serial transfer was observed with *A. flavus* (45). Together with our work on *Penicillium*, these studies demonstrate the rapid and coordinated phenotypic and metabolomic shifts in a range of filamentous fungi as they are stably maintained in high-resource environments.

One of the most striking changes that we observed in our *Penicillium* mutants is a shift in VOC production. Even though the domestication process in our experiments was undirected, white mutants stopped producing the musty VOC geosmin that is generally considered undesirable in foods (31) and increased production of ketones and other VOCs that are considered desirable in Camembert and other cheeses (11, 32). The genes and pathways responsible for production of the secondary metabolite geosmin and other VOCs are not well-characterized in cheese *Penicillium* species (46) so we are unable to explicitly link the transcriptomic data with the VOC data. Ketone production by cheese fungi is a result of lipases releasing fatty acids from lipids that are then converted into ketones, alcohols, and other VOCs (25, 46, 47). The observed shifts in VOC production could reflect a generalized loss of secondary metabolite production and increased lipid degradation of the cheese substrate.

We did not identify a specific genetic mechanism controlling the observed trait evolution in *Penicillium* and a genetic mechanism underlying degeneration of *Aspergillus* cultures has also not been identified. It is possible that genomic evolution, including structural changes such as chromosomal rearrangements or mutations in specific genes, could drive the phenotypic and metabolomic shifts observed. Alternatively, the rapid trait evolution in *Penicillium* may be explained not by genomic evolution, but by transgenerational epigenetic inheritance which has been proposed to be important in filamentous fungi (48). Global regulators of genes involved with pigment, toxin, and spore production have been identified in *Aspergillus* and other fungi and some of these regulators, including the methyltransferase LaeA, have been demonstrated to epigenetically regulate transcription (49, 50). Future work further characterizing these mutants will identify the specific genetic and molecular changes driving domestication.

A detailed record of how contemporary *P. camemberti* mutants used in cheese production were derived is not available (13), so we cannot know precisely how and when *P. commune* was domesticated to become *P. camemberti*. It is possible that the industrial starter cultures used today were isolated as mutants from cheese caves in Europe. Regardless of how these mutants were ultimately acquired, our work demonstrates the potential for *Penicillium* molds to rapidly evolve without intentional selection for desired cheese-making traits. Because we observed similar trait shifts in two different *Penicillium* species, it is possible that phenotypic mutants of many different *Penicillium* species are continuously evolving in cheese caves around the world. Our laboratory domestication suggests that new strains of *Penicillium* for cheese production could be generated through more intentional and controlled domestication processes. Most strains used in mold-ripened cheese production originate from Europe, providing a limited palette of textures and flavors. The experimental domestication of *Penicillium* on cheese could lead to the production of locally adapted strains for cheese production in other parts of the world.

## METHODS

### Isolation and manipulation of *Penicillium* cultures

Two non-starter *Penicillium* strains, *Penicillium* sp. 162_3FA and *Penicillium* sp. 12, were used in the experiments throughout this study. Both fungal strains were isolated from the surface of a natural rind cheese produced and aged in Vermont, USA. A third non-starter mold, *Penicillium* sp. MB was isolated from a natural rind cheese production and aging facility in California. It was sequenced as part of this work and included in the phylogenomic analysis, but was not used in the experimental portion of the paper because mutant scoring was not as clear as with the other two strains. To determine the putative taxonomic identity of these two molds, whole genome sequences were obtained using Illumina sequencing as previously described for a *Mucor* isolate (51). Genomes were assembled using the *de novo* assembler in CLC Genomics Workbench and annotated using GenSAS (https://www.gensas.org/).

### Construction of a phylogenomic tree of *Penicillium*

To reconstruct the evolutionary relationships among described *Penicillium* species and the two isolates used in this work, we obtained a comprehensive set of genomes from *Penicillium* species using NCBI’s Taxonomy Browser. We downloaded the 33 available *Penicillium* genomes on February 5^th^ 2018. In addition to the 33 *Penicillium* genomes, we also downloaded three genomes from representative species in the genus *Aspergillus* for use as outgroup taxa. Altogether, our dataset contains a total of 39 taxa – three *Penicillium* genomes sequenced in the present study, 33 publicly available *Penicillium* genomes, and three *Aspergillus* taxa.

To identify orthologous genes, we used the Benchmarking Universal Single-Copy Orthologs (BUSCO), version 2.0.1 (52), pipeline and the Pezizomycotina database (creation date: 02-13-2016) from ORTHODB, version 9 (53). The BUSCO pipeline uses hidden Markov models for 3,156 previously established Pezizomycotina orthologs (hereafter referred to as BUSCO genes) to individually search each genome for their corresponding orthologs. Identified orthologs are classified as “single copy” if only a single full length orthologous gene is identified, “duplicated” if two or more full length predicted orthologs are identified, “fragmented” if the identified orthologous gene is less than 95% of the sequence length of the gene’s multiple sequence alignment used for constructing the hidden Markov model, or “missing” if no orthologous gene is found in the genome.

Examination of BUSCO genes reveals that the genomes sequenced and assembled in the present study exhibited high genome completeness. For example, using 3,156 universally single-copy orthologs from Pezizomycotina (or BUSCO genes), we observed that the MB_DraftGenome, *Penicillium commune* (162_2_DraftGenome), and 12_DraftGenome isolates have 3,099 / 3,156 (98.2%), 3,097 / 3,156 (98.2%), 3,096 / 3,156 (98.1%) of BUSCO genes present in their genomes, respectively. Notably, these values are similar to those of other publically available genomes; for example, *P. citrinum* and *P. camemberti* have 3,095 / 3,156 (98.1%) and 3,100 / 3,156 (98.3%) of BUSCO genes present in their genomes, respectively.

To construct the phylogenomic data matrix, we first retained only those BUSCO genes that were present in single copy in at least 20 taxa (i.e., > 50% taxon-occupancy). From this set of 3,111 groups of BUSCO orthologous genes, we created individual gene alignments using MAFFT, version 7.249b (54)(Katoh and Standley, 2013) with the BLOSUM62 matrix of substitutions (Mount, 2008), the “geneafpair” parameter, a gap penalty of 1.0, and 1,000 maximum iterations. To thread nucleotide sequences on top of each amino acid multiple sequence alignment, we used a custom PYTHON, version 3.5.2 (https://www.python.org/), script using BIOPYTHON, version 1.7. Individual nucleotide alignments were trimmed using TRIMAL, version 1.4 (55), with the “automated1”. Thereafter, the 3,111 groups of BUSCO orthologous genes were concatenated into a single phylogenomic data matrix that contained 5,498,894 sites and had taxon occupancy of 97.65 ± 5.23%.

To reconstruct the evolutionary history of *Penicillium* species, we used the gene-based maximum likelihood schemes of concatenation and coalescence (56–58). We first determined the best-fitting models of sequence substitutions and conducted maximum likelihood searches of single gene trees for all 3,111 genes in our data matrix. Using IQ-TREE, version 1.6.1 (59), we determined the best-fitting model for each single gene or partition according to the Bayesian Information Criterion. We inferred each gene’s evolutionary history using the “-m TEST” parameter and distinct randomly generated seeds specified using the “-seed” parameter. Among best-fitting models, the most commonly observed one was “TN+F+I+G4” (60, 61), which was observed in 537 / 3,111 (17.26%) genes, “TIM2+F+I+G4” (61), which was observed in 440 / 3,111 (14.14%) genes, and “GTR+F+I+G4”, which was observed in 400 / 3,111 (12.86%) genes (61, 62).

To infer evolutionary history using the gene-based concatenation approach, we first created a partition-file that describes the best-fitting parameters for each gene and conducted five independent searches for the maximum likelihood topology. More specifically, to allow each of the 3,111 genes in the phylogenomic data matrix to have its own substitution model and rate heterogeneity across sites parameters, we created a nexus-format partition file that detailed these parameters and gene boundaries. Using this file along with the “-spp” parameter, each partition was also allowed to have its own set of evolutionary rates (63). We also increased the number of candidate trees used during the search for the optimal tree from 5 to 10 using the “-nbest” parameter. Using this scheme, we conducted 5 independent searches with 5 distinct seeds for the maximum likelihood topology. We next evaluated support for the maximum likelihood topology using the ultrafast bootstrap approximation approach (UFBoot) (64).

To infer evolutionary history using coalescence, we combined all single gene phylogenies into a single file. Using this file, we inferred the evolutionary history of *Penicillium* species using ASTRAL-II, version 4.10.12 (57) using default parameters and assessed bipartition support using local posterior probability (65).

Examination of the resulting phylogeny revealed high concordance with previous whole-genome based analyses (66) with full support at every internode using UFBoot and local posterior probability except for the split of section *Exilicaulis* and *Lanata-divaricata*, which received local posterior probability value of 0.97. Importantly, our analyses place the newly sequenced *Penicillium commune* in section of *Fasciculata* (Fig A), as previously described (67).

To explore putative biological drivers of incongruence in the phylogenies, we determined how many individual gene trees supported any or none of the three alternative topologies by calculating their gene support frequencies (68). More specifically, we counted how many single gene trees supported a sister group relationship of *Penicillium biforme* and *Penicillium camemberti,* how many a sister group relationship of *P. biforme* and *Penicillium commune*, and how many a sister group relationship of *P. camemberti* and *P. commune*. We also counted how many gene trees did not infer these three species as a monophyletic group. Finally, we also kept track of how many genes could not be used due to incomplete taxon representation to determine if incomplete taxon representation was contributing to incongruence. Gene support frequencies were determined by examining the sister species of *Penicillium commune* using NEWICK UTILITIES, version 1.6 (69).

### Experimental evolution of *Penicillium* on cheese

Each of the *Penicillium* strains was grown in our experimental cheese system consisting of 20 mL of cheese curd agar (CCA) in a standard 100 x 15 mm Petri dish. Cheese curd agar (CCA) is composed of freeze-dried unsalted cheese curd from a blue cheese produced in Vermont (100 g/L), xanthan gum (5 g/L), salt (30 g/L) and agar (17 g/L). CCA allows for controlled manipulations of cheese rind communities and accurately mimics the dynamics of cheese rind development (14). To start the evolution experiment, each strain was initially inoculated with 500 colony-forming units (CFUs) across the surface of the CCA plate. Each experimental cheese community was incubated for 7 days in the dark at 24°C and 95% humidity. Experimental communities were serially transferred to new CCA every week for a period of 8 weeks.

To manipulate the biotic environment throughout the evolution experiment, cheese rind bacteria and yeasts were added to four replicate communities to create a “*Penicillium* + Community” treatment. The yeast *Debaryomyces hansenii* strain 135B and the bacteria *Staphylococcus xylosus* BC10 and *Brachybacterium alimentarium* strain JB7 were added at the same density as *Penicillium* at the initial inoculation. We selected these three microbial species for the *Penicillium* + Community treatment because they represent taxa that are common members of natural rind cheese microbiomes (14, 16, 51) and were stably maintained during the duration of the experiment. We acknowledge that these community members may have evolved during the experimental domestication experiment and we did not attempt to control for their evolution throughout the experiment.

To manipulate total resource availability throughout the evolution experiment, we created “low cheese” which contained the same components as CCA except 10 g/L of freeze-dried unsalted cheese curd (instead of 100 g/L in “normal cheese”) was used in the medium. The pH of “low cheese” was identical to that of “normal cheese”. In the “alternating normal/low” treatment, we alternated transfers each week between full-strength and dilute CCA, starting with full-strength CCA when setting up the experiment.

At each transfer, the cheese curd agar from each community was removed from the Petri dish and homogenized inside a Whirl-Pak bag containing 30 mL of 1X phosphate buffered saline (PBS). From this homogenized mixture, an aliquot of 100 µL was plated onto new CCA to seed a new community. Another aliquot was serially diluted and plated onto PCAMS media with 50 mg/L of chloramphenicol (to inhibit bacterial neighbors in the “*Penicillium* + Community” treatment) for colony counting and scoring colony phenotypes. Glycerol stocks were made at each transfer so that communities could be archived and revived later if needed.

Phenotypic evolution was tracked by scoring wild-type and mutant colonies at each transfer. Mutant colonies were considered to have differences in pigment intensity, distribution of pigment around the colony, colony texture, degree of sporulation, and extent of mycelium production (explained in detail in Table S1A-S1B). Phenotyping was done after 5 days of incubation of PCAMS plates containing the output of each transfer. PCAMS output plates were incubated at 24°C for 5 days before phenotyping was completed. Phenotyping occured on plates with at least 100 colonies.

### Reproductive and mycotoxin trait analysis

Reproductive and mycotoxin traits were only measured for the ancestor and select mutants of *Penicillium* sp. 162_3FA because it is most closely related to *P. camemberti*. The following strains were used in these assays: Ancestor, M2, M3, M5, M6, M7, M9, and M10. These strains were selected for trait profiling because they spanned the spectrum of visible colony types ranging from similar to wild-type (slightly less blue) to completely white (Table S1A-S1B). To determine reproductive output, each strain was inoculated on three replicate plates at a density of 50 CFUs on the surface of 20 mL of cheese curd agar (CCA) in a standard 100 x 15 mm Petri dish. At this density, individual CFUs were discernible. Plates were incubated for 7 days at 24°C and 95% humidity. From three individual colonies, a sterile circular cork borer with a diameter of 0.7cm was inserted into the center of the colony. The excised colony plug was serially diluted and CFUs were determined on PCAMS.

To determine how production of mycotoxin cyclopiazonic acid changed in mutant strains compared to ancestors, we measured CPA production in the following strains: Ancestor, M2, M3, M5, M6, M7, M9, and M10. 40,000 CFUs of each strain was spread across the surface of 20 mL cheese curd in a 100 x 15 mm Petri dish. Three biological replicates of each strain were used in the experiments. Plates were incubated in the dark for 3 days at 24°C and 11 days at 4°C. After the 14 day incubation, the medium was harvested from the plate, placed into a Whirl-pak bag, and homogenized. Samples were frozen at −80°C until analysis.

The cyclopiazonic acid concentration of the cheese curd agar was measured using liquid chromatography with tandem mass spectrometry (LC-MS/MS) at Romer Labs (Union, Missouri, USA). The homogenized cheese curd sample was extracted in a 50/50 mixture of acetonitrile and deionized water by shaking for 90 minutes. The supernatant was filtered and 10 mL was mixed with 500 µL of acetic acid. 1 mL of this solution was vortexed in a MycoSpin Column (Romer Labs), vortexed for 1 minute, and centrifuged for 30 seconds at 10,000 rpm. 75 µL of the purified extracted was injected into a Shimadzu HPCL with a Phenomenox Gemini HPCL C18 column (4.6 X 150 mm, 5 µm) with mobile phase A consisting of ESI + 5 mM ammonium formate with 0.1% formic acid in deionized water and mobile phase B consisting of acetonitrile. The injection volume was 40 µL, the flow rate was 1.0 mL/min, and the column temperature was 40°C. Internal standards of CPA were used to construct a calibration curve.

Stability of traits was assessed in two mutant strains M5 and M6. Three replicate plugs of each of these strains was transferred to fresh CCA weekly using heat-sterilized stainless steel cork borers (6mm diameter). Colonies were photographed at each transfer as in **Fig. S5**.

### Competition experiments

We competed ancestor *Penicillium* sp. 162_3FA with the mutant strain M9 to determine whether evolved mutant strains have a higher fitness compared to the ancestor. It is challenging to standardize input densities of the white mutants because they produced many fewer spore compared to the ancestor. Mutant M9 of *Penicillium* sp. 162_3FA was chosen as a competitor because it still produced significant numbers of spores making it possible to make comparable initial inoculum for both the ancestor and evolved strains. Experiments were conducted in 96-well plates with 150µL of 10% CCA added to each well and 200 CFUs of each strain added at the start of the experiment. Six replicate experimental cheese communities containing the wild-type and mutant mix were incubated in the dark at 24°C for ten days. To determine the abundance of WT and M9 at the end of the experiment, each replicate community was removed from the 96-well plate, homogenized in 600 µL 1X PBS, serially diluted onto PCAMS, and then WT and M9 colonies were counted.

### Volatile profiling

Cheese volatiles were collected from fungal cultures by headspace sorptive extraction (HSSE) using a glass encapsulated magnetic stir-bar coated with 0.5mm polydimethylsiloxane (PDMS). Before each sample was collected, the stir-bars were heated from 40°C to 300°C at 5°C/min and flushed with 50 mL/min nitrogen (Airgas) to desorb sorbed organics using a Gerstel (Baltimore, MD) TC2 tube conditioner. HSSE is an equilibrium-driven, enrichment technique in which 10mm long stir-bars, Twister^TM^ from Gerstel, were suspended 1 cm above the sample by placing a magnet on the top side of the collection vessel cover. All cultures were sampled in quadruplicate (n=4) for 4 hr. One replicate of the Control (WT) was lost during sample processing. After collection, the stir-bar was removed and spiked with 10 ppm ethylbenzene-d_10_, an internal standard obtained from RESTEK (Bellefonte, PA). Organics were introduced into the gas chromatograph/mass spectrometer (GC/MS) by thermal desorption. In addition to Twister blanks, analysis of the agar media was made to ensure background interferents were minimal. If present, these compounds were subtracted from the fungal data.

Analyses were performed using an Agilent (Santa Clara, CA) 7890A/5975C GC/MS equipped with a 30 m x 250 µm x 0.25 µm HP5-MS column. The instrument was equipped with an automated multi-purpose sampler (MPS), thermal desorption unit (TDU), and a CIS4 programmable temperature vaporizing (PTV) inlet from Gerstel. The TDU, operating in splitless mode, transferred the sample from the stir-bar to the CIS4, which was held at −100°C, by ramping the temperature from 40°C to 275°C at 720°C/min, then held isothermal for 3 min under 50 ml/min helium gas flow. Once transferred, the CIS4 was heated from −100°C to 280°C at 12°C/min, and held for 5 min. The GC temperature was held at 40°C for 1 min, then ramped to 280 °C at 5°C/min, and held for 5 min. The MS was scanned from 40 to 250 *m/z*, with the EI source at 70 eV. A standard mixture of C7 to C30 n-alkanes, purchased from Sigma–Aldrich (St. Louis, MO), was used to calculate the retention index (RI) of each compound in the sample.

The Ion Analytics (Gerstel) spectral deconvolution software was used to analyze the GC/MS data. Peak identification was performed through comparison of sample and reference compound spectral patterns and retention indices, NIST05, Adams Essential Oil Library, and literature. Compound identity was based on the following set of conditions. First, peak scans must be constant for five or more consecutive scans (differences ≤ 20%). Second, the scan-to-scan variance (SSV or relative error) must be < 5. The SSV calculates relative error by comparing the mass spectrum at each peak scan against another. The smaller the difference, the closer the SSV is to zero, the better the MS agreement. Third, the Q-value must be ≥ 93. The Q-value is an integer between 1 and 100; it measures the total ratio deviation of the absolute value of the expected minus observed ion ratios divided by the expected ion ratio times 100 for each ion across the peak. The closer the value is to 100, the higher the certainty between database and sample spectra. Finally, the Q-ratio compares the ratio of the molecular ion intensity to confirming ion intensities across the peak; it also must be ≤ 20%. When all criteria are met, the software assigns a compound name or numerical identifier to the peak from the database.

To cluster the VOC data, a UPGMA tree with 100 bootstraps was constructed using a Bray-Curtis dissimilarity matrix in PAST3. Analyses of similarity (ANOSIM) was used to test whether there were differences between the ancestor and mutant in VOC profiles. ANOSIM R values indicate the degree to which groups separate, with 1 being complete separation and 0 indicating a complete lack of separation. Similarity percentage analysis (SIMPER) on Bray-Curtis dissimilarity distances was used to identify the compounds that contributed most to differences in VOC profiles.

### RNA-sequencing

To determine global changes in gene expression in cheese-adapted *Penicillium*, we compared the ancestor and mutant M5 of *Penicillium* sp. 162_3FA were inoculated onto cheese curd agar. Inoculum of both strains came from streaks of the fungi growing on plate count agar with milk and salt (PCAMS) that had been growing for one week. A 1 cm^2^ plug was taken from the leading edge of mycelium and then homogenized in 500 µL of phosphate buffered saline (PBS). At three evenly spaced locations on a 100cm wide Petri dish containing 20 mL of cheese curd agar, 20 µL of the inoculum was spotted onto the agar surface. After 72 hours of growth in the dark at 24 °C, the spots were 1.5cm in width. The wild-type 162_3FA had produced spores and was blue in color and the mutant M5 162_3FA was white in color. The entire fungal mass from each of the three spots was cut away from the CCA and then placed in RNAlater (Qiagen) and stored at −80 °C. Four biological replicates were sampled for each of the two strains.

RNA was extracted from one of the three spots from each replicate plate using a Qiagen RNeasy Plant Mini Kit after grinding the sample in liquid nitrogen with an autoclaved mortar and pestle. Approximately 100 mg of ground fungal biomass was placed in 750μl of Buffer RLT with 10μl of β-mercaptoethanol per 1 ml added to the Buffer RLT. The manufacturer’s recommended protocol was followed for RNA extraction, including an on column DNAse treatment. To isolate mRNA, the NEBNext ® Poly (A) mRNA Magnetic Isolation Module (New England Biolabs) was used. This mRNA was used to generate RNA-seq libraries using the NEBNext ® Ultra II RNA Library Prep Kit for Illumina following the manufacturer’s recommended protocol. The RNA-seq libraries were sequenced using 125 base-pair length, paired-end Illumina sequencing on a HiSeq at the Harvard Bauer Core.

After trimming low quality sequences and removing failed reads using CLC Genomics Workbench, sequencing yielded 3.5 to 22 million forward reads that were used for read mapping and differential expression analysis. Reads were mapped to a reference genome of *P. commune* 162_3FA that was sequenced using pair-end 125 base-pair length Illumina sequencing, assembled with CLC Genomic Workbench de novo assembler, and annotated using GenSAS (https://www.gensas.org/). Read mapping was performed with the CLC Genomics Workbench RNA-seq analysis pipeline with the following settings: mismatch cost of 2, insertion cost of 3, deletion cost of 3, length fraction of 0.8, and similarity fraction of 0.8. The number of unique reads mapped (mapped to one specific gene and not additional locations in the genome) was used to determine expression levels and quantile normalization was used to take into account different levels of sequencing across replicates. Other methods of calculating gene expression and normalization were assessed (e.g. RPKM) and did not change the main findings of the differential expression analysis. Identification of genes that were differentially expressed in the cheese-adapted mutant compared to the ancestor was completed by using the empirical analysis of differential gene expression tool in CLC Genomics Workbench. This pipeline uses the exact test for two-group comparisons (70). We considered those genes with greater than 5-fold change in expression and FDR corrected *p*-values of < 0.05 as differentially expressed genes. To identify specific biological pathways that were enriched in the sets of downregulated or upregulated genes, we used a KOBAS 2.0 to conduct a hypergeometric test on functional assignments from the gene ontology (GO) database (using the *Aspergillus flavus* genome as a reference for GO ID assignment) with Benjamini and Hochberg FDR correction.

### Cheese cave population population sampling

Sterile toothpicks were used to sample rinds of 43 wheels of a natural rind blue cheese in the same caves where *Penicillium* sp. Strain 12 had been previously isolated. Samples were placed in 1X PBS, stored at 4°C for 24 hours, and then each sample from a wheel of cheese was plated onto PCAMS with chloramphenicol to inhibit bacterial growth. Plates were incubated at 24°C for seven days before assessing plates for the presence of white mutants. Camembert-style cheeses inoculated with *P. camemberti* are aged in different caves at the same facility. To confirm that white mutant phenotypes were derivatives of the wild-type *Penicillium* sp. strain 12 and not contamination from starter cultures, we used whole genome sequencing as described above to sequence a white mutant morphotype isolate. Read mapping using BowTie2 revealed that the genome had a 99.9% pairwise identity to the reference genome of *Penicillium* sp. strain 12.

### Data availability

Whole-genome sequences of *Penicillium commune* strain 162_3FA and *Penicillium* sp. strain 12 have been submitted to NCBI as MUGJ00000000 and MUGI00000000, respectively. Raw data from RNA-sequencing of *Penicillium* sp. 162_3FA strain Ancestor and *Penicillium* sp. 162_3FA strain M5 and resequencing the cave isolate of *Penicillium* #12 sp. have been deposited in the NCBI Sequence Read Archive as PRJNA510622.

## ACKNOWLEDGEMENTS

This work was supported by the National Science Foundation (1715553). Freddy Lee, Esther Miller, Casey Cosetta, Pat Kearns, and Megan Biango-Daniels provided very helpful feedback on an earlier version of this manuscript.

## SUPPLEMENTAL FIGURE and TABLE LEGENDS

**Figure S1: The genome-scale phylogeny of the genus *Penicillium***

(A) Concatenation phylogeny with section denominations. The two strains used in the experiments described in the text (shown in bold) are placed within section *Fasciculata*. Furthermore, *Penicillium commune* strain 162_3FA is closely related to *Penicillium biforme* and *Penicillium camemberti*. Inset depicts phylogeny with branch lengths representing substitutions per site. *Penicillium* sp. MB was isolated from a natural rind cheese at the same time as the other two strains and was sequenced as part of this work, but it was not used in the experiments described. (B) Comparison of concatenation-based (left) and coalescence-based (right) phylogenies reveals only one instance of incongruence. Specifically, whereas *P. biforme* is placed sister to *P. commune* 162_3FA in the concatenation analysis, coalescence supports *P. camemberti* as sister to *P. commune* 162_3FA. All internal branches received full support except the coalescence-inferred internal branch where *Exilicaulis* and *Lanata-divaricata* split, which received a local posterior probability value of 0.97. Branch lengths reflect substitutions / site for concatenation and coalescence units for the coalescence inferred phylogeny.

**Figure S2: Population size of *Penicillium commune* 162_3FA when evolved alone and with a community of cheese microbes.** Lines connect points representing mean colony forming units (CFUs) of four replicate populations and error bars represent one standard deviation of the mean. Total CFUs in the *Penicillium* + community treatment was significantly different from *Penicillium* alone (repeated-measures ANOVA *F*_1,6_= 10.3, *p* = 0.02).

**Figure S3: Experimental evolution of *Penicillium* sp. 12 alone and with a community of cheese rind microbes. (A)** Population size of *Penicillium* sp. 12 when evolved alone and with a community of cheese microbes. Lines connect points representing mean mutant phenotype frequencies of four replicate populations and error bars represent one standard deviation of the mean. Total CFUs in the *Penicillium* + community treatment was significantly different from *Penicillium* alone (repeated-measures ANOVA *F*_1,6_= 16.8, *p* =0.006). Lines connect points representing mean colony forming units (CFUs) of four replicate populations and error bars represent one standard deviation of the mean. **(B)** Phenotypic mutant frequency of *Penicillium* sp. 12 when evolved alone and with a community of cheese microbes. Lines connect points representing mean mutant phenotype frequencies of four replicate populations and error bars represent one standard deviation of the mean. Mutant frequency in the *Penicillium* + community treatment was significantly different from *Penicillium* alone (repeated-measures ANOVA *F*_1,6_= 20.1, *p* <0.005).

**Figure S4: Population size of *Penicillium commune* 162_3FA when evolved in different cheese nutrient environments.** “Normal cheese” = 10% cheese curd in agar medium. “Low cheese” = 1% cheese curd in agar medium. “Alternating normal/low” = alternating 10% and 1% cheese curd at each transfer. The “Low cheese” treatment suppressed population size (repeated-measures ANOVA *F*_2,9_= 105.1, *p* <0.0001, with Tukey’s HSD post-hoc tests). Lines connect points representing mean colony forming units (CFUs) of four replicate populations and error bars represent one standard deviation of the mean.

**Figure S5: Stability of *Penicillium commune* 162_3FA mutant phenotypes.** Mutants were transferred weekly to new cheese curd agar and colony morphology was photographed. The white mutant morphology was stable over time.

**Table S1A:** Description of mutant classes and specific mutant isolates identified in the evolution of *Penicillium commune* strain 162_3FA.

**Table S1B:** Distribution of mutant types across replicate populations and transfers in the experimental evolution of *Penicillium* commune 162_3FA. Data below show the number of colonies of Ancestor or mutants counted from 10-4 dilution plates of experimental populations. T1, T2, etc. = transfer number.

**Table S2A:** Description of specific mutant isolates identified in the evolution of *Penicillium* sp. strain #12

**Table S2B:** Distribution of mutant types across replicate populations and transfers in the experimental evolution of *Penicillium* sp. #12. Data below show the number of colonies of Ancestor or mutants counted from 10-4 dilution plates of experimental populations. T1, T2, etc. = transfer number.

**Table S3:** Overview of all volatile organic compounds detected in the ancestor (ANC), and mutants M2, M5, and M6 of *Penicillium commune* strain 162_3FA.

**Table S4:** Overview of genes that were differentially expressed bewteen the ancestor and mutant M5 of *Penicillium commune* strain 162_3FA

