## Supplementary material for "Rapid phenotypic and metabolomic domestication of wild *Penicillium* molds on cheese": Figure S1

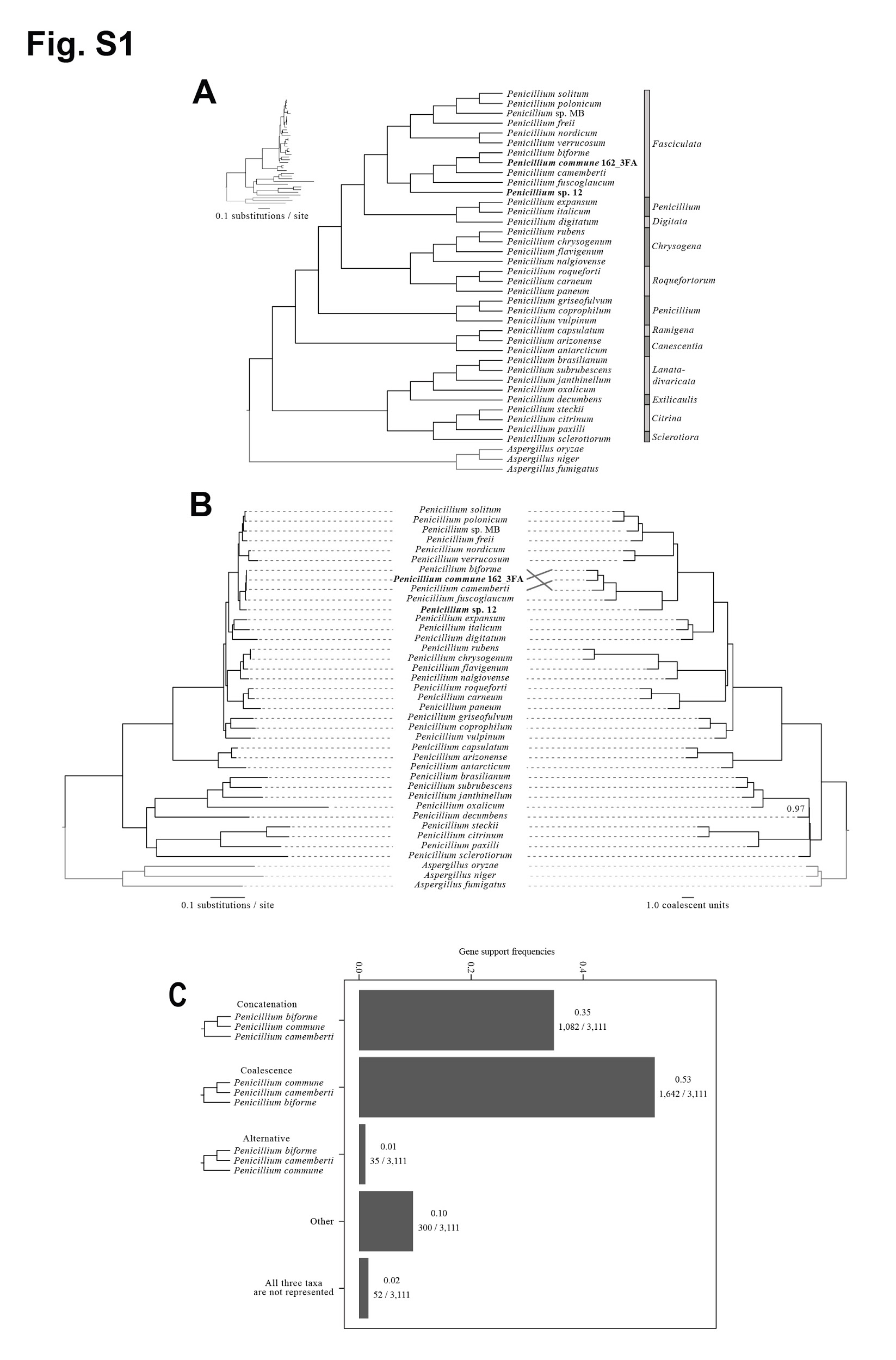

**Figure S1: The genome-scale phylogeny of the genus *Penicillium***

(A) Concatenation phylogeny with section denominations. The two strains used in the experiments described in the text (shown in bold) are placed within section *Fasciculata*. Furthermore, *Penicillium commune* strain 162_3FA is closely related to *Penicillium biforme* and *Penicillium camemberti*. Inset depicts phylogeny with branch lengths representing substitutions per site. *Penicillium* sp. MB was isolated from a natural rind cheese at the same time as the other two strains and was sequenced as part of this work, but it was not used in the experiments described. (B) Comparison of concatenation-based (left) and coalescence-based (right) phylogenies reveals only one instance of incongruence. Specifically, whereas *P. biforme* is placed sister to *P. commune* 162_3FA in the concatenation analysis, coalescence supports *P. camemberti* as sister to *P. commune* 162_3FA*.* All internal branches received full support except the coalescence-inferred internal branch where *Exilicaulis* and *Lanata-divaricata* split, which received a local posterior probability value of 0.97. Branch lengths reflect substitutions / site for concatenation and coalescence units for the coalescence inferred phylogeny.
