## Supplementary material for "Rapid phenotypic and metabolomic domestication of wild *Penicillium* molds on cheese": Figure S2

**Figure S2: Population size of *Penicillium commune* 162_3FA when evolved alone and with a community of cheese microbes.** Lines connect points representing mean colony forming units (CFUs) of four replicate populations and error bars represent one standard deviation of the mean. Total CFUs in the *Penicillium* + community treatment was significantly different from *Penicillium* alone (repeated-measures ANOVA *F*_1,6_= 10.3, *p* = 0.02).
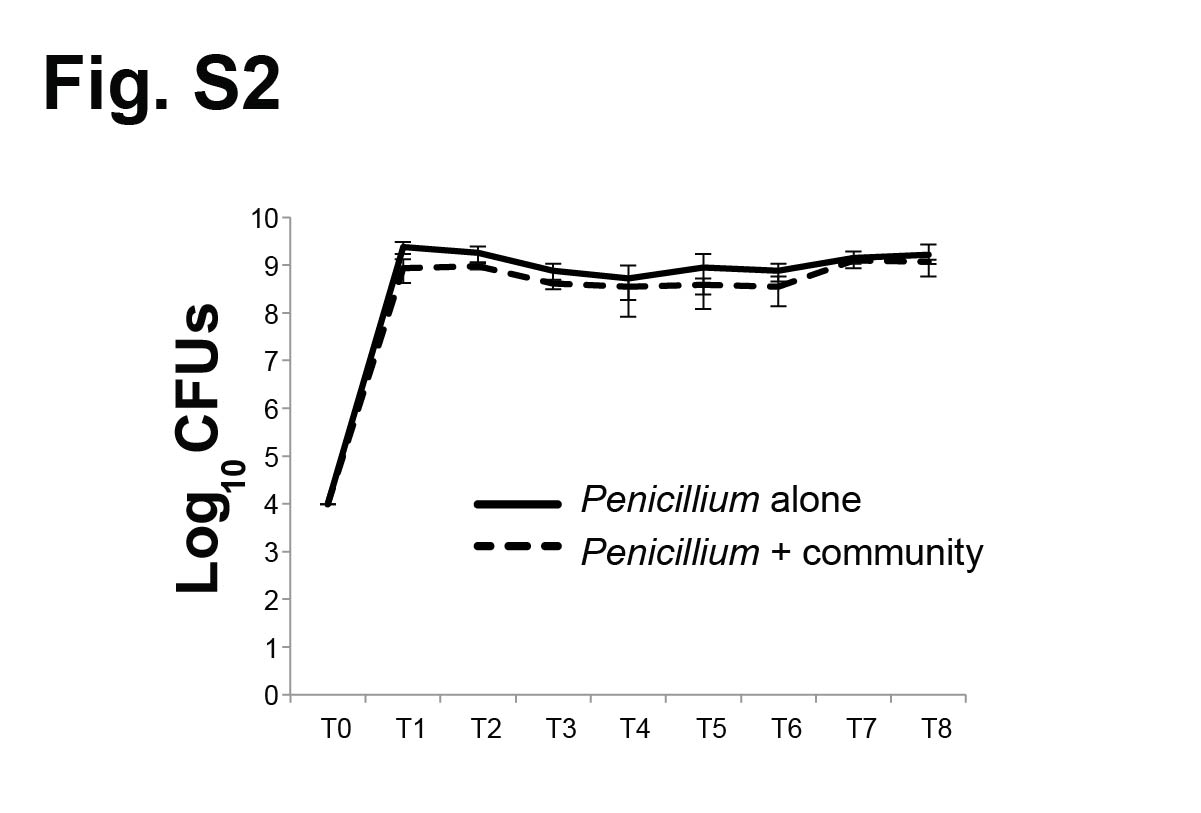
