## Supplementary material for "Rapid phenotypic and metabolomic domestication of wild *Penicillium* molds on cheese": Figure S3

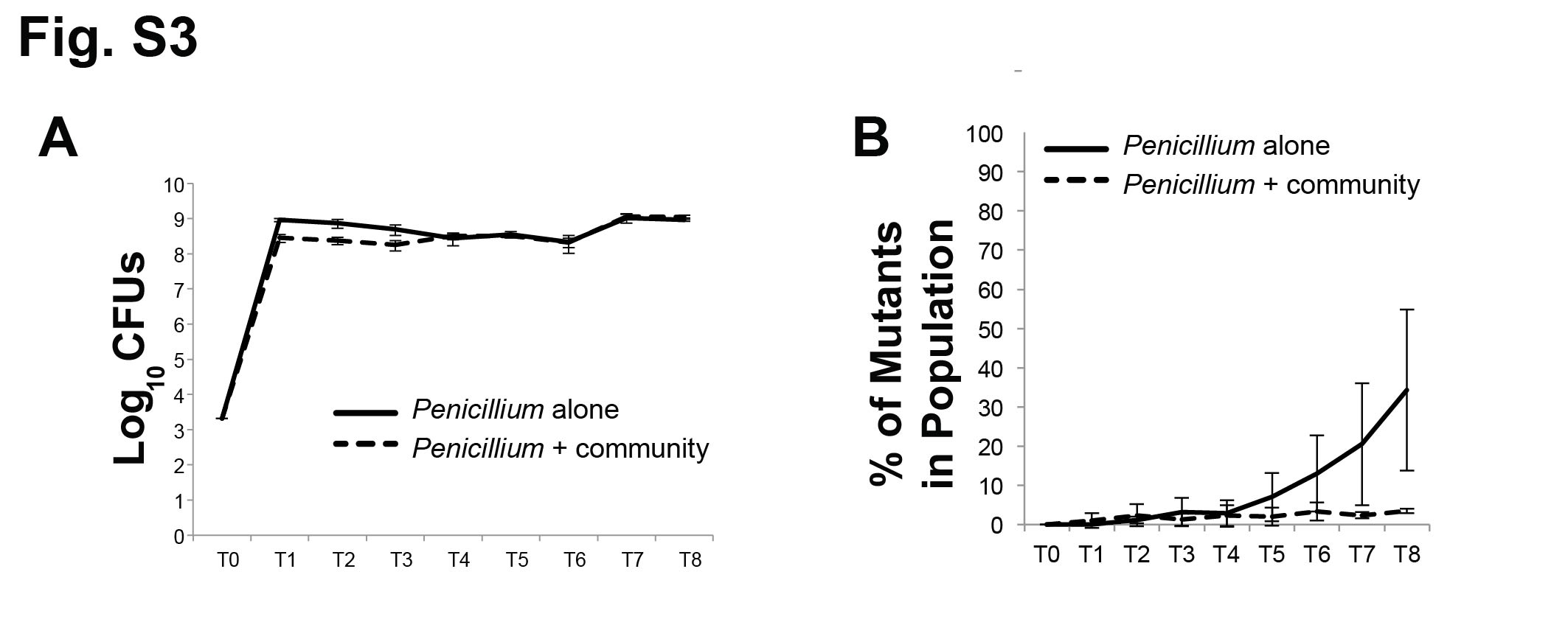

**Figure S3: Experimental evolution of *Penicillium* sp. 12 alone and with a community of cheese rind microbes. (A)** Population size of *Penicillium* sp. 12 when evolved alone and with a community of cheese microbes. Lines connect points representing mean mutant phenotype frequencies of four replicate populations and error bars represent one standard deviation of the mean. Total CFUs in the *Penicillium* + community treatment was significantly different from *Penicillium* alone (repeated-measures ANOVA *F*_1,6_= 16.8, *p* =0.006). Lines connect points representing mean colony forming units (CFUs) of four replicate populations and error bars represent one standard deviation of the mean. **(B)** Phenotypic mutant frequency of *Penicillium* sp. 12 when evolved alone and with a community of cheese microbes. Lines connect points representing mean mutant phenotype frequencies of four replicate populations and error bars represent one standard deviation of the mean. Mutant frequency in the *Penicillium* + community treatment was significantly different from *Penicillium* alone (repeated-measures ANOVA *F*_1,6_= 20.1, *p* <0.005).
