## Supplementary material for "Rapid phenotypic and metabolomic domestication of wild *Penicillium* molds on cheese": Figure S4

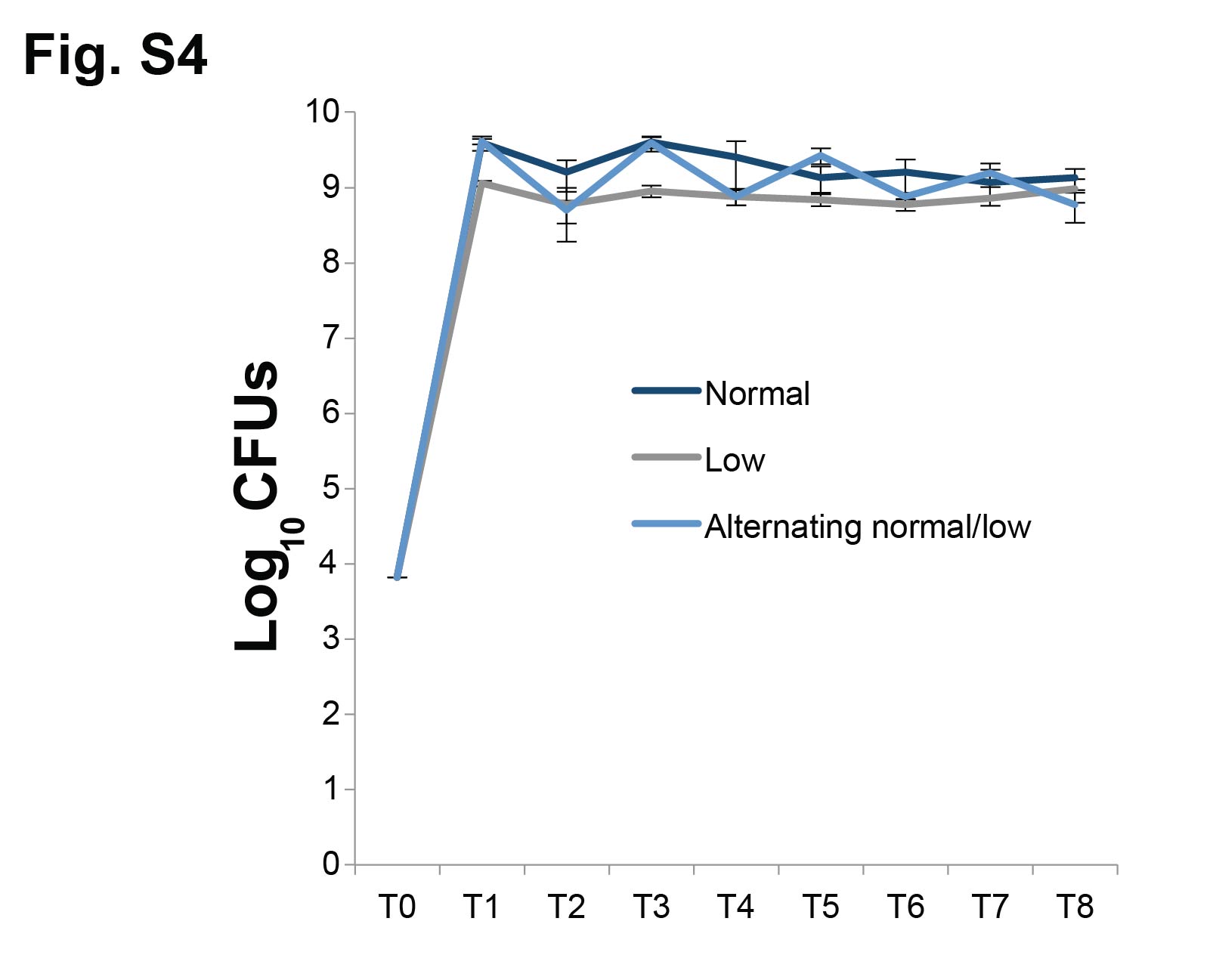

**Figure S4: Population size of *Penicillium commune* 162_3FA when evolved in different cheese nutrient environments.** “Normal cheese” = 10% cheese curd in agar medium. “Low cheese” = 1% cheese curd in agar medium. “Alternating normal/low” = alternating 10% and 1% cheese curd at each transfer. The “Low cheese” treatment suppressed population size (repeated-measures ANOVA *F*_2,9_= 105.1, *p* <0.0001, with Tukey’s HSD post-hoc tests). Lines connect points representing mean colony forming units (CFUs) of four replicate populations and error bars represent one standard deviation of the mean.
