## Supplementary material for "Rapid phenotypic and metabolomic domestication of wild *Penicillium* molds on cheese": Figure S5

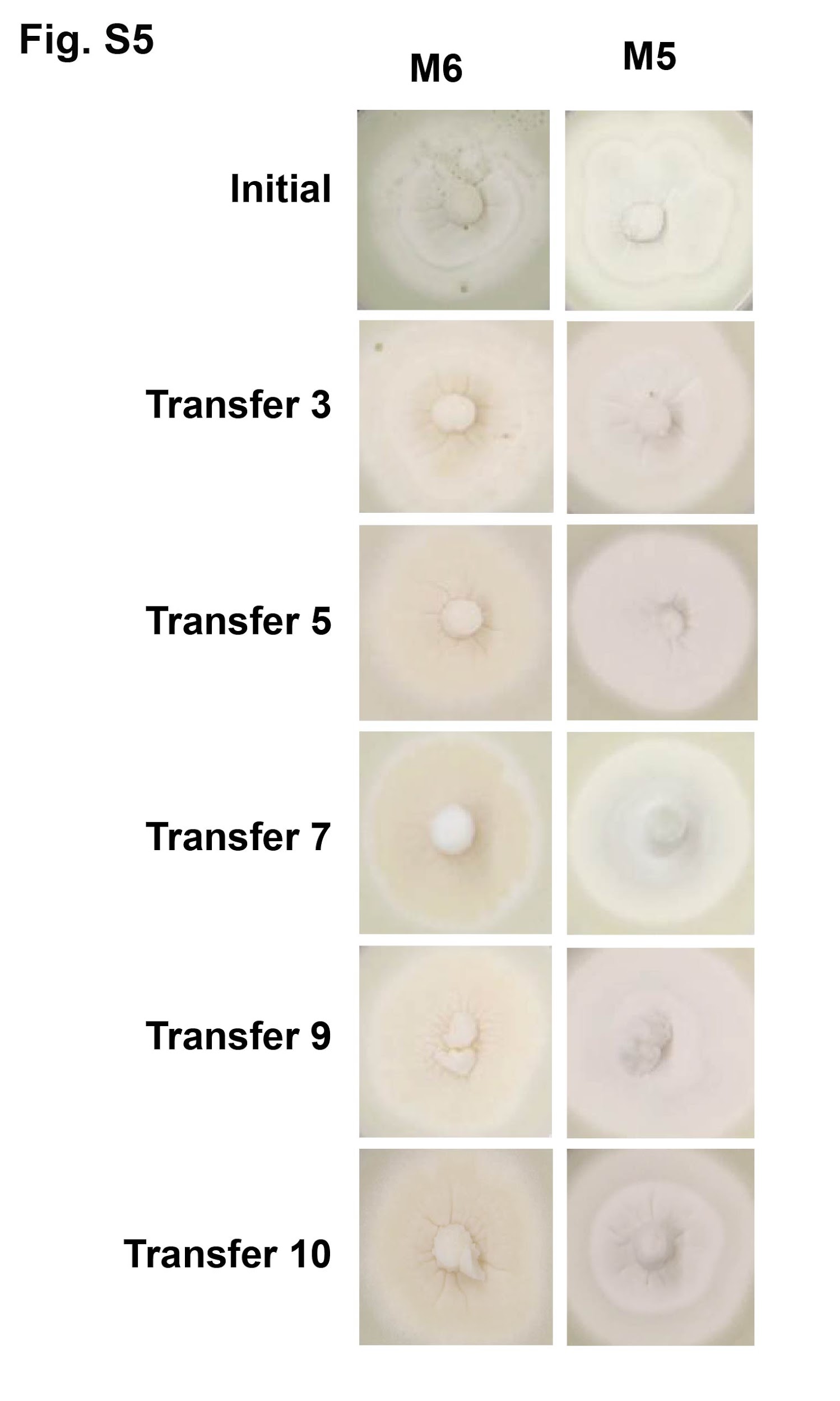

**Figure S5: Stability of *Penicillium commune* 162_3FA mutant phenotypes.** Mutants were transferred weekly to new cheese curd agar and colony morphology was photographed. The white mutant morphology was stable over time.
